# Shifts in competitive structures can drive variation in species’ phenology

**DOI:** 10.1101/2022.12.13.520270

**Authors:** Patricia Kaye T. Dumandan, Glenda M. Yenni, S. K. Morgan Ernest

## Abstract

For many species, a well-documented response to anthropogenic climate change is a shift in various aspects of its life-history, including its timing or phenology. Often, these phenological shifts are associated with changes in abiotic factors used as proxies for resource availability or other suitable conditions. Resource availability, however, can also be impacted by competition, but the impact of competition on phenology is less studied than abiotic drivers. We fit generalized additive models (GAMs) to a long-term experimental dataset on small mammals monitored in the southwestern United States and show that altered competitive landscapes can drive shifts in breeding timing and prevalence, and that, relative to a dominant competitor, other species exhibit less specific responses to environmental factors. These results suggest that plasticity of phenological responses, that is often described in the context of annual variation in abiotic factors, can occur in response to biotic context as well. Variation in phenological responses under different biotic conditions shown here further demonstrates that a more nuanced understanding of shifting biotic interactions is useful to better understand and predict biodiversity patterns in a changing world.

## INTRODUCTION

Characterizing species’ responses to various environmental conditions is useful in determining their sensitivity and adaptive capacity to the consequences of ongoing global environmental change (Glick et al., 2011). Among the known responses to climate change, shifts in phenology or the timing of key life-history events of organisms, are the most well-documented (Cohen et al., 2018; Roslin et al., 2021; Visser & Both, 2005; Walther et al., 2002). Often, the timing of important phenological events (e.g., reproduction) match with periods of high resource availability to support high fitness and survival costs (Bradshaw & Holzapfel, 2007; Buss et al., 2021; Forchhammer et al., 1998; Helm et al., 2013; Reed et al., 2013; Walther et al., 2002). Fitness costs of reproduction are often attributed to the increased energetic demands of maintaining and developing reproductive organs and behaviors (Audzijonyte & Richards, 2018) and survival costs are incurred from increased predation risk (Kelt et al., 1995). To avoid unfavorable conditions during the reproductive period, species may use abiotic (i.e., temperature, day length, precipitation) and biotic cues (i.e., flowering, prey availability) to regulate and shape reproductive phenology (Denny et al., 2014; Kenagy & Bartholomew, 1981; Wolkovich et al., 2014).

Timing reproductive activities to overlap with periods of high resource availability is important for consumer species because decoupled resource and consumer phenology can negatively impact production and survival of offspring (Thackeray et al., 2010; Wann et al., 2019). Thus, many studies assess phenological drivers related to either reliable environmental cues of resource availability or biotic cues related directly to the emergence or abundance of prey (i.e., resource) species (DeLucia et al., 2012; Gordo & Sanz, 2005; Pearce-Higgins & Yalden, 2004). However, resource availability is determined by more than just the intrinsic population dynamics of prey. Competition from other individuals can deplete important resources and create resource limitation even when resource supply is initially high. Recent theoretical models (Carter & Rudolf 2022) have demonstrated that resource depletion from competition (intra- or inter-specific) favors early initiators of reproduction over latter initiators and drives high phenological synchrony as a result.

Understanding how competition impacts phenology is important because, as species enter and leave communities in response to environmental changes, the timing and magnitude of resource availability perceived by resident species may also change (Carter & Rudolf 2022) and promote shifts in both timing and the apparent response to abiotic cues. If phenology is fixed to specific abiotic cues or if competition is a process of lesser importance, then we would expect removing competitors to have little impact on the timing or magnitude of reproductive events. We would also see little difference in the important drivers associated with reproductive activity under different competitive scenarios. However, changes to reproductive phenology in response to competition are possible, given its potential to alter resource availability. The study by Carter and Rudolf (2022) posits that competitive impacts can differ depending on whether a focal species initiates their phenology concurrent to or before their competitor. Late initiation allows a dominant competitor to draw down valuable resources, reducing reproductive success for the focal species. Moreover, assuming plasticity in an individual’s ability to decide to enter a reproductive state, the presence of a competitor may drive decreases in the prevalence of reproductive individuals. If reproductive timing is a fixed trait, only individuals with certain phenological timings that reduce the competitive impact may enter breeding condition. However, while species’ interactions (i.e., competition) can theoretically impact reproductive timing and investment, empirical assessment of these impacts are limited.

Long-term natural field experiments, where competitive networks are explicitly manipulated, are well-suited for investigating the role of intra- and inter-specific competition in driving population processes, including reproduction (Ernest et al., 2016; Kelt et al., 1995, Mysterud et al., 2008; Ostfeld, 1985). In long-term field experiments where dominant competitors are removed, the effect of their loss on the reproductive phenology of other species in the system can be evaluated by comparing the reproductive parameters of a focal species in plots with and without a dominant competitor. Here, we used long-term data on small mammals monitored in an experimental field site near Portal, Arizona to assess the effects of competition on reproductive phenology. This long-term experiment (1977 to present) in the southwestern USA generates fine-scale (monthly) data on desert rodents under different biotic contexts by experimentally removing a dominant competitor, the kangaroo rats (*Dipodomys* spp.) in a subset of the experimental plots. In this study, we compared the breeding phenology of desert pocket mice (*Chaetodipus penicillatus*) and Bailey’s pocket mice (*Chaetodipus baileyi*) in *Dipodomys* accessible (or Dipo*+*) and *Dipodomys* inaccessible (or Dipo-) plots to determine the effects of interspecific competition on the phenology of natural populations. Specifically, we determined periods of significant change in the monthly proportions of breeding male and female individuals in both plot types and assessed the sex-specific responses to environmental conditions, and intra- and inter-specific competition of both species in each plot type (Dipo*+* and Dipo-).

## METHODS

### Study System

To assess the impacts of competition on reproductive phenology, we used data from a long-term field experiment located in the Chihuahuan Desert near Portal, Arizona (31°56’20.29”N 109° 4’47.44”W). Initiated in 1977, this study site consists of 24 experimental plots (50m x 50 m) divided up among treatments that manipulate the desert rodent community. Each plot is fenced, and we manipulate access to plots using different sized openings through the fencing to generate some plots containing an unmanipulated rodent community (Dipo+), a community where *Dipodomys* spp. (three species of behaviorally dominant competitors of which *Dipodomys merriami* is the most common) are only occasionally present in low numbers (Dipo-), and plots where all rodents are excluded (total rodent exclosure). In this study, we only used data from the Dipo+ and Dipo- plots. Because of the spatial extent of the experiment (the plots are contained in a 20-ha area), the rodents share the same habitat and experience the same environment, allowing us to assess the impacts of competition without correlated changes in abiotic conditions or habitat structure. All plots were sampled monthly, and the sex, species, and reproductive condition of each individual were recorded affording us a monthly record of reproductive activity for every species in the community. Treatments on some plots have changed over time, so for this study we restricted our analysis to data collected from 1988 to 2014 (Appendix S1: Fig. 1), the longest period when the experimental treatments were consistent for the most plots. More detailed description of the experimental plots is discussed elsewhere (Ernest et al. 2016).

**Figure 1.**
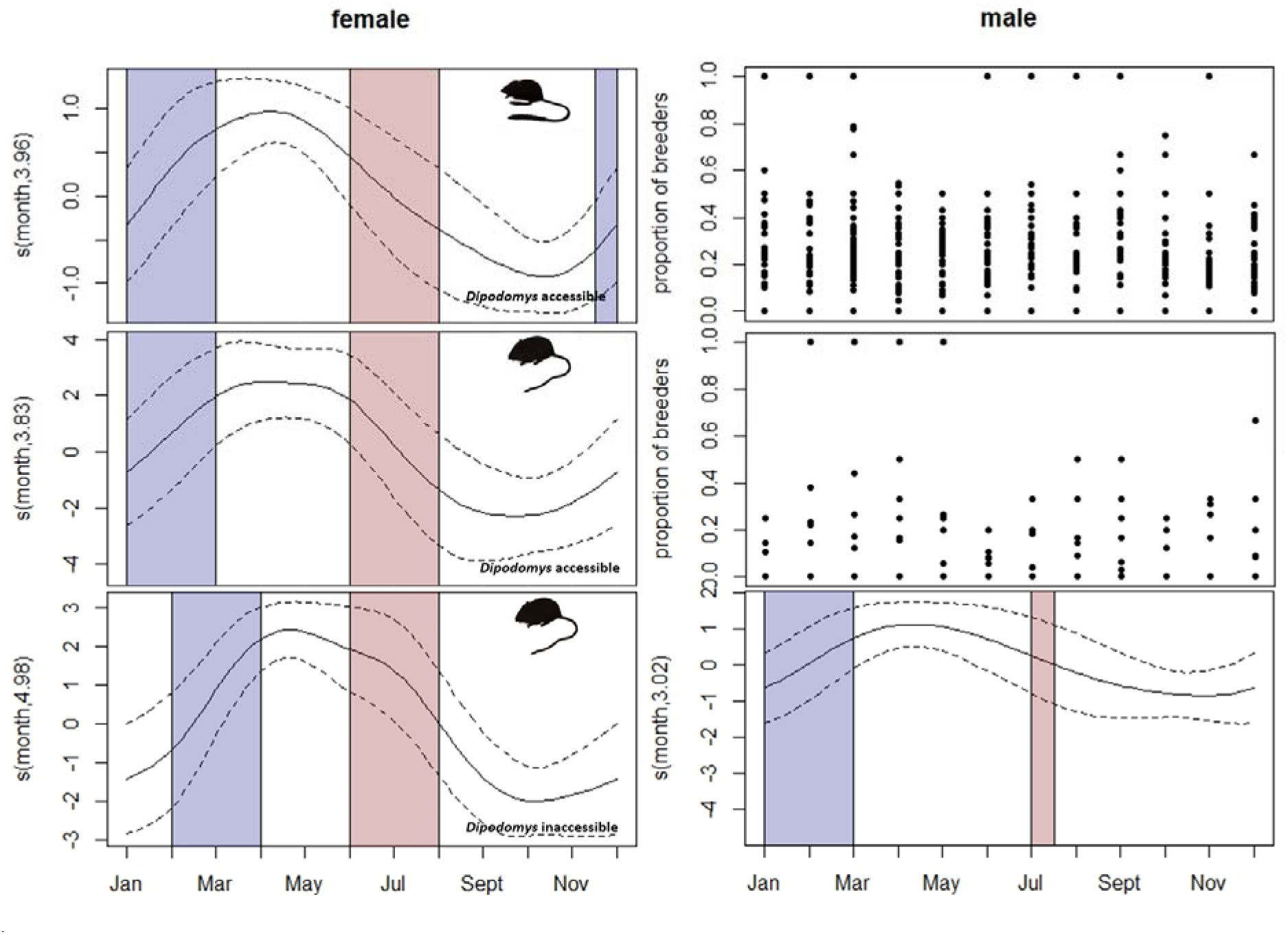
Reproductive phenology of female (left panel) and male (right panel) *Dipodomys* spp. in *Dipodomys* accessible plots (top panel), and *C. baileyi* in *Dipodomys* accessible (middle panel) and inaccessible (bottom panel) plots in an experimental field site near Portal, Arizona. Plots show estimated smooths of the month term from species-, sex-, and treatment-specific generalized additive models (GAMs). Blue and red shading indicates periods of significant increase and decrease in breeding proportions, respectively. In cases where seasonal patterns or periods of significant increase or decrease could not be estimated from the model, raw data on proportion of breeding individuals were plotted (male *Dipodomys* spp. and *C. baileyi*;).

### Rodent data

Desert rodents are an ideal model system for asking questions about the role of competition on phenology because male and female reproductive investments are largely known in rodents (Mysterud et al., 2008) and competition is a strong structuring process in desert rodent communities (Brown and Munger 1985, Bledsoe and Ernest 2019). We focused our analyses on two species, *C. baileyi* and *C. penicillatus*, because they are numerous on both target plot types between 1988-2014 and because previous work has demonstrated strong competitive interactions between each other and with *Dipodomys* spp. (Bledsoe and Ernest 2019). Because the timing of phenology initiation relative to the dominant competitor is important in Clark and Rudolf’s experiment (2022), we also analyzed the temporal dynamics of *Dipodomys*’ breeding for comparison. We used the ‘portalr’ package (Christensen et al., 2019) to extract a monthly time series by species (*Dipodomys* spp., *C. baileyi*, *C. penicillatus*), treatment (Dipo+, Dipo-), and sex (male, female) of the abundance (i.e., all individuals regardless of reproductive status) and counts of only reproductive individuals. Males were considered in reproductive state if they were recorded as scrotal (descended), recently scrotal, or had minor signs of scrotal testes. Females were considered reproductive if recorded as pregnant (after palpating the belly), and/or exhibited either red and/or enlarged nipples, and/or those with swollen and/or plugged vaginas. To limit the introduction of biases in our dataset through the inclusion of juveniles, we identified a minimum threshold of the body mass for the breeding individuals (McLean and Guralnick, 2020) for each species and sex (i.e., *C. baileyi* male= 16 g, *C. baileyi* female= 21 g, *C. penicillatus* male= 10 g, *C. penicillatus* female= 12 g).

### Environmental data

We used monthly data on NDVI (an index of plant greenness), mean temperature (°C), cumulative precipitation (millimeters; sliding window calculating rainfall accumulations) that fell during warm or cool months (calculated as the sum of precipitation on days when minimum temperature is > or < 4 °C, a biologically relevant cutoff for the transition between warm and cool rainy seasons which generate different plant assemblages), and biomass data of *C. baileyi*, *C. penicillatus*, and all *Dipodomys* spp. We also examined lags of 0 and 1 month, and cumulative values (i.e., precipitation) for environmental data because weather affects vegetation and population dynamics dynamically through temporal processes (Ding et al., 2020; Wen et al., 2019) and the importance of lags and seasonal accumulations have repeatedly been demonstrated in explaining rodent population dynamics at this site (Thibault et al 2010). We used biomass data of *C. baileyi*, *C. penicillatus*, and all *Dipodomys* spp. as proxies for competitive effects (henceforth referred to as biotic factors).

### Data Analysis

We built generalized additive models (GAMs) to characterize the reproductive phenology of male and female *C. baileyi* and *C. penicillatus* in different treatments, and to determine the relationship between the proportion of breeding individuals with environmental (i.e., weather and resource availability) and biotic (i.e., the intraspecific and interspecific competitive context) factors. We also built GAMs to characterize the reproductive phenology of *Dipodomys* spp. to compare any shifts in the breeding phenology of *Chaetodipus* spp. with that of the dominant competitor. We pooled all data on *Dipodomys* spp. monitored at the site (*D. ordii*, *D. merriami*, *D. spectabilis*) and focused only on their timing in Dipo+ plots, where we have adequate observation data on the group. We used GAMs because they are appropriate for fitting data that may exhibit non-linear trends and associations (Pedersen et al. 2019), a dynamic typical of species’ phenology that wax and wane through time.

#### Quantifying between-treatment differences in phenology

To assess the difference in the breeding phenology of rodents on Dipo+ and Dipo- plots, we estimated a separate smoother for month, and for each plot type, which were treated as ordered factors (Rose et al. 2012). By treating the treatments as ordered factors (P_0_= Dipo+, P_1_= Dipo-), we were able to determine difference in seasonal trends (i.e., smooths) between the Dipo + (i.e., treated as the reference level) and Dipo- plots. We built species- and sex-specific models that had the general form:

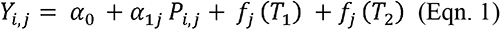

where Y is the response variable (i.e., proportion of breeding individuals)*, α*_0_ is the model intercept (i.e., the mean value of *Y* in the reference treatment, *P*_0_; i.e., Dipo+ plot), *α*_1_ is the difference in the mean response between the Dipo+ (*P*_0_) and Dipo-plot (*P*_1_), *f*() are smooth functions of month (*T*_1_), representing the breeding phenology of rodents, that are parameterized using cubic cyclic regression splines (Pederson et al. 2019), and of year (*T*_2_), representing long term trends, that are parameterized using Gaussian process smooths (Wood 2017) for the *j*th treatment.

#### Fitting the drivers of breeding phenology

To assess how the competitive environment may impact model fits and the relative importance of drivers, we built species-, sex- and treatment-specific models that included terms for environmental factors (terms for NDVI, warm and cool precipitation, and mean monthly temperature with lag of zero and one), a proxy for intraspecific competition (i.e., population biomass), and for interspecific competition (i.e., biomass of competitors) as biotic (i.e., competitive context) factors. Pairwise correlation tests showed low correlation among environmental covariates used to fit all models (*r* < |0.5|; see Appendix for results). We built separate models for each treatment, species, and sex combination. These models followed the general form:

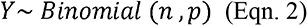

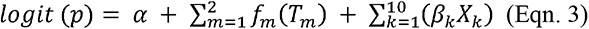

In Eqn. 2, Y is the number of breeding individuals, which follows a binomial distribution with the parameters *n* (number of independent trials, e.g., total number of males) and *p* (probability of success, e.g., proportion of breeding males). In Eqn. 3, *p* is modeled as a function of the intercept (*α*), the smooth function for month (*f*_l_(*T*_l_)), the smooth function for year (*f*_2_(*T*_2_)), and linear terms for the effects of environmental and biotic predictors (*β_k_*). Similar parameterizations of the smooth terms done in Eqn. 1 were followed. We standardized all predictors by centering each value on the mean and dividing them by two standard deviations (Gelman, 2008) to facilitate interpretation of results. All models were implemented in a restricted maximum likelihood (REML) framework using the ‘mgcv’ package ver 1.8-31 (Wood 2017) in R ver 4.1.1 (R Core Team, 2021). Data (<u>10.5281/zenodo.7301687</u>) and code (<u>10.5281/zenodo.8121230</u>) used to conduct data analyses are archived on Zenodo.

### Model Interpretation

#### Reproductive phenology

From Eqn. 1, we determined significant treatment effects and visualized differences between treatments by plotting the difference smooths, which reflect the estimated differences in seasonality between Dipo+ and Dipo- plots. Then, we determined the beginning and end of each breeding season for each species-, sex-, treatment-combination by identifying periods of significant change in the proportion of breeding individuals for each category using the output from models built using Eqn. 3. Here, the first derivatives of the seasonal spline (*f*_l_(*T*_l_)) were estimated using finite differences to find the time periods when the derivatives are significantly increasing or decreasing (i.e., does not contain zero in the confidence interval). This and the 95% pointwise confidence interval were determined using the derivatives function in the ‘gratia’ package (Simpson, 2023). We interpreted periods when the rate of change in the seasonal spline (*f*_l_(*T*_l_)) were increasing as the onset of breeding season (i.e., when more individuals are exhibiting reproductive characteristics), and decreasing rates of change as the termination of breeding season (i.e., when fewer individuals are exhibiting reproductive characteristics). We also performed exploratory analyses on the reproductive phenology of *C. penicillatus* at different time periods of *C. baileyi* establishment in the site. *C. baileyi* colonized the site around 1995, causing major shifts in resource use and the competitive landscape, especially for *C. penicillatus* (Bledsoe and Ernest 2019, Diaz and Ernest 2022). To assess the effect of the shift in the competitive landscape caused by this natural experiment, we also built simpler models (without terms for competition) for all species-, sex-, and treatment-specific datasets to determine the relative importance of these terms in explaining variation in breeding proportions. Methods and results of these exploratory works are in the Appendix.

#### Association between covariates and proportion of breeders

We assessed whether individuals of both pocket mice species exhibited different responses to drivers with and without *Dipodomys* spp. present. We interpreted the coefficient values based on their signs (positive or negative) to assess whether the effect of a given factor on the proportion of breeding individuals was positive or negative. We also used the distribution curves of the estimated slopes from the models to better understand how *Chaetodipus* spp. phenology relates to its dominant competitor.

## RESULTS

Both *Chaetodipus* spp. exhibited changes in their breeding patterns when *Dipodomys* spp. were removed from the rodent community. Changes included shifts in breeding onset (i.e., *C. baileyi* females and to some extent, males), and prevalence of breeding individuals (*C. penicillatus* females). Some relationships between covariates and the proportion of breeders did not differ between *Dipodomys* accessible and *Dipodomys* inaccessible plots. However, the responses of both *Chaetodipus* spp. to most covariates were broader than that of *Dipodomys* spp. in Dipo+ plots and were narrower in Dipo- plots.

### Female Bailey’s pocket mouse (C. baileyi)

Female *C. baileyi* on Dipo+ plots went into breeding condition earlier than those on Dipo - plots (*s*(difference)= 1.55, p= 0.05, deviance explained= 46.7%; January to March vs. February to April, respectively; Fig.1, Appendix S1: Fig. S2), but declines in reproductive activity occurred during similar timeframes on both Dipo+ and Dipo- plots (June-August vs. July-Sept, respectively; Fig. 1). On both plots, the onset of the breeding season for female *C. baileyi* occurred later than that for females of the dominant competitor, *Dipodomys* spp. (Fig. 1). Thus, in the absence of *Dipodomys* spp., female *C. baileyi* delayed reproduction by ∼1 month and in the presence of *Dipodomys* spp., entered reproductive condition earlier and stayed in reproductive condition longer.

The best fit models for female *C. baileyi* on Dipo+ and Dipo- inaccessible plots overlapped in their important drivers, but also contained important differences. On Dipo+ plots, female reproductive condition was positively associated with warm precipitation (lag 0; amount of warm precipitation on corresponding month) and cool precipitation (lag 1; amount of cool precipitation a month prior; variation explained= 66.4%; R^2^= 0.57; Fig. 2) while on Dipo- plots, warm precipitation (lag 0) was still significant but cool precipitation was not. Additional important associations that emerged in the Dipo- model that did not appear in the Dipo+ model included a negative association with NDVI (lag 0), a positive association with NDVI (lag 1), and a positive association with *C. penicillatus* biomass (variation explained= 69.3%; R^2^= 0.66; Fig. 3). Distributions of the covariate slope estimates (Fig. 4 and Appendix S1: Fig. S5) exhibited broader distributions on Dipo+ plots for some environmental and biotic factors than either *Dipodomys* spp. or *C. baileyi* on Dipo- plots.

**Figure 2.**
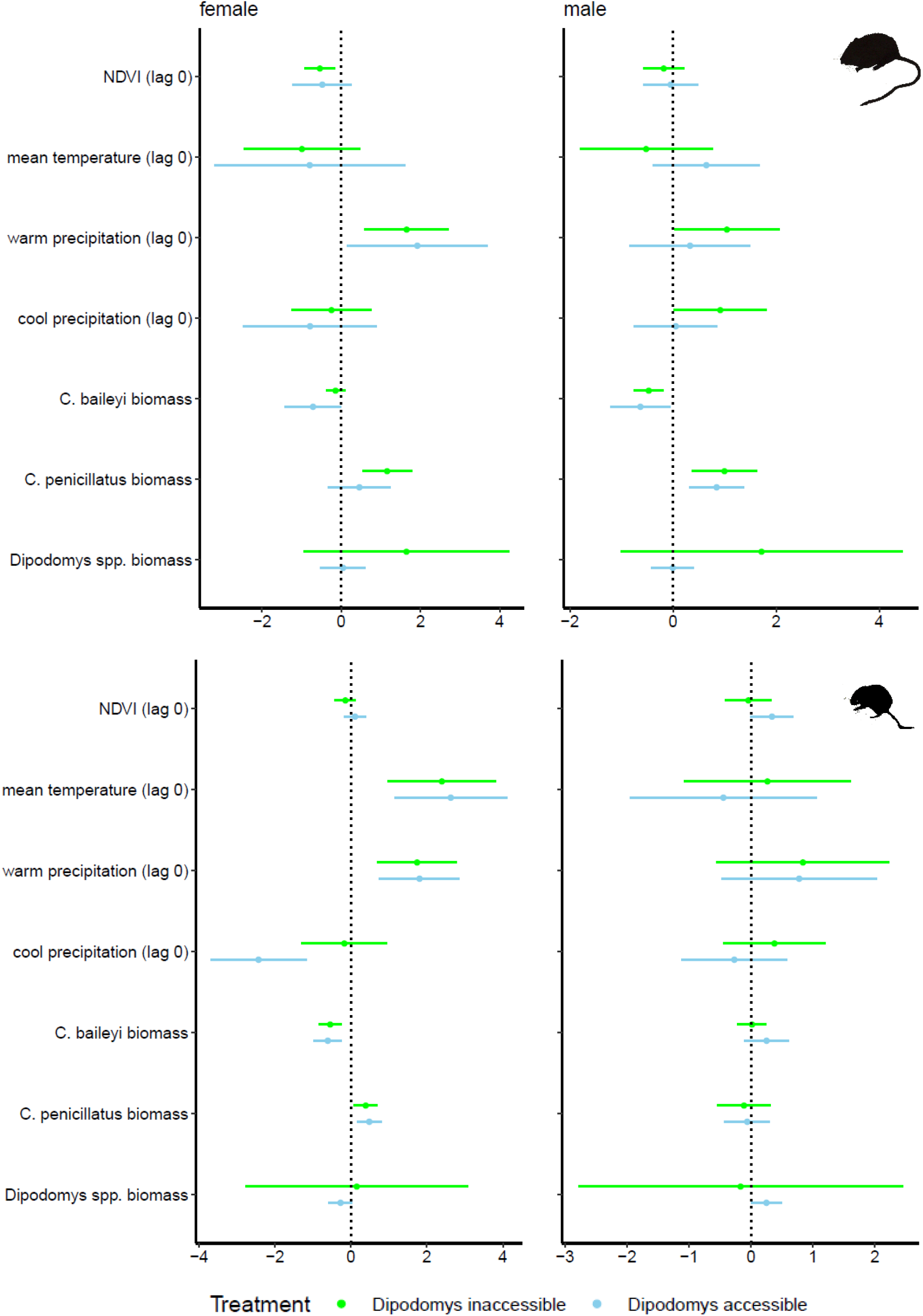
Association between select environmental and biotic factors with the proportion of reproductive female (left panel) and male (right panel) *C. baileyi* (top panel) and desert pocket mouse (bottom panel) in *Dipodomys* accessible and inaccessible plots.

**Figure 3.**
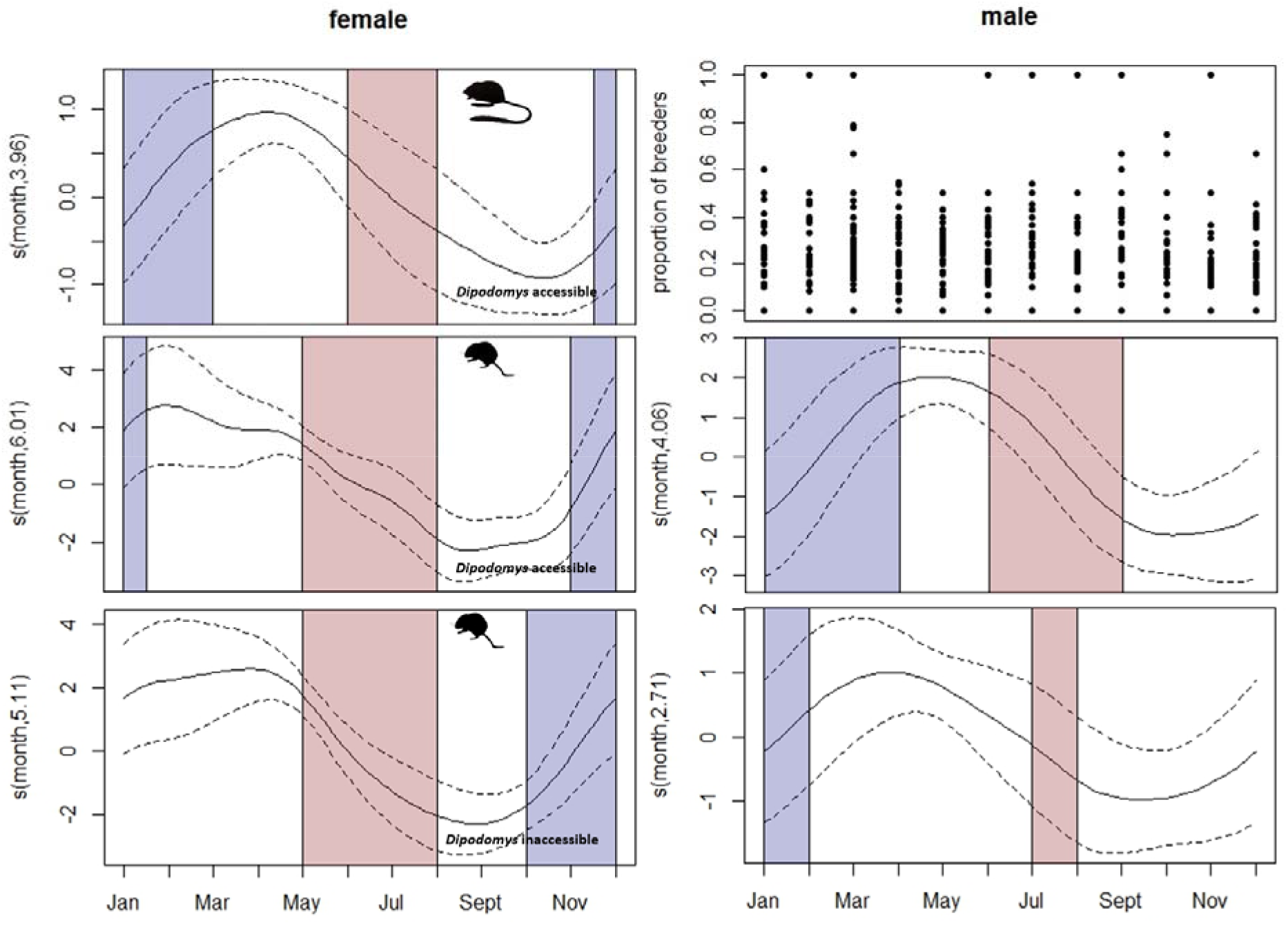
Reproductive phenology of female (left panel) and male (right panel) *Dipodomys* spp. in *Dipodomys* accessible plots (top panel), and *C. penicillatus* in *Dipodomys* accessible (middle panel) and inaccessible (bottom panel) plots in an experimental field site near Portal, Arizona. Details on the configuration of these plots are like Figure 1.

**Figure 4.**
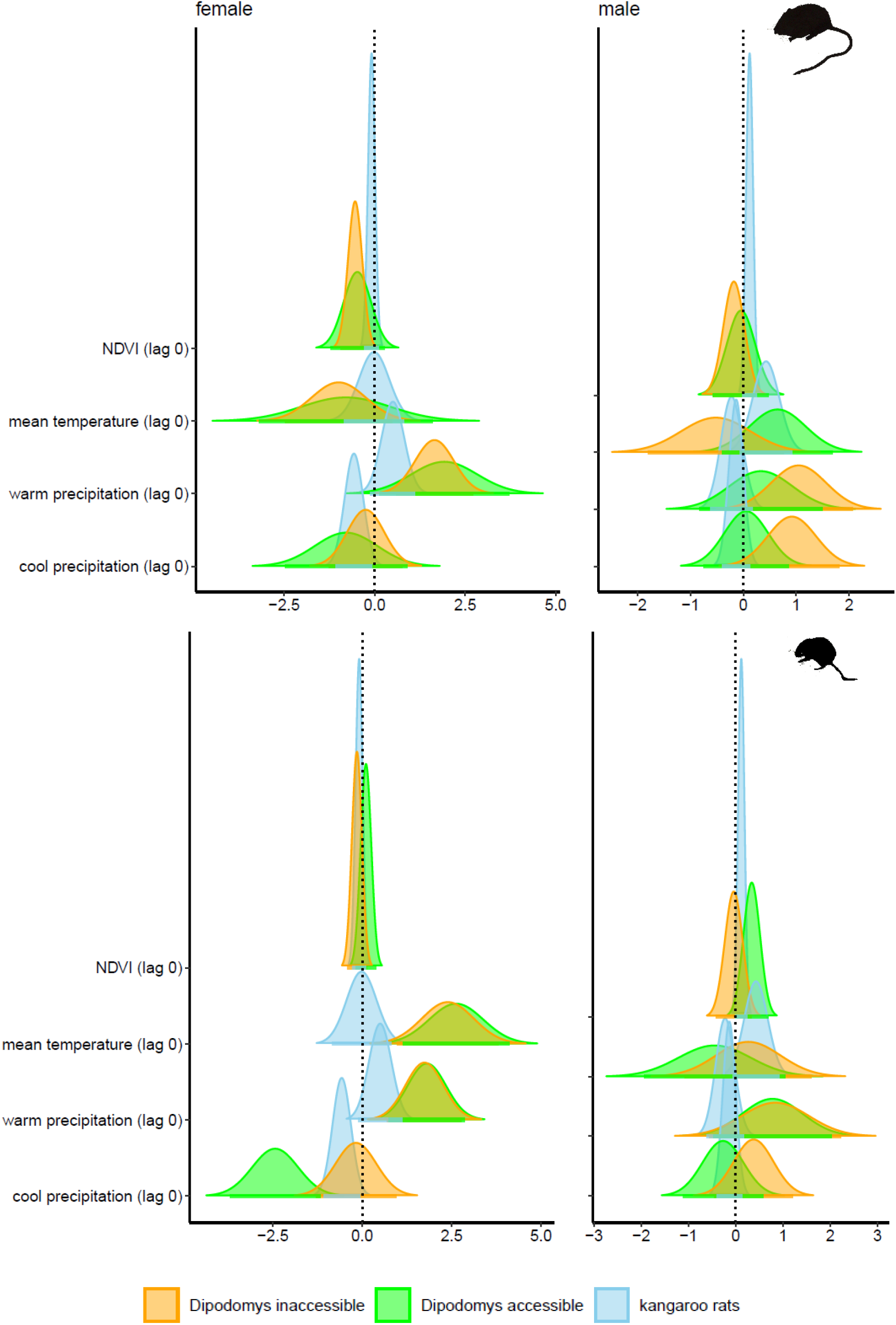
Distribution curves of the abiotic covariate slope estimates indicating the response of female (left panel) and male (right panel) *C. baileyi* (top panel) and *C. penicillatus* (bottom panel) in *Dipodomys* acessible and inaccessible plots, compared to *Dipodomys* spp. estimates. Curves indicate the normal distribution with mean equal to the point estimate, and standard deviation equal to the standard error of the coefficients.

### Male Bailey’s pocket mouse (C. baileyi)

The breeding timing of male *C. baileyi* did not significantly differ between Dipo+ and Dipo- plots (*s*(difference= 1.35, *p*= 0.11, deviance explained= 20.7%; Appendix S1: Fig. S2). However, despite a lack of a statistically significant treatment effect, the treatments exhibit a slight divergence in their seasonal smooths that peak around June-July. The environmental GAMs fit separately to Dipo+ and Dipo- plots were also suggestive of a difference in the seasonal signal – with the Dipo+ model exhibiting no seasonal signal whereas a weak seasonal signal was detected on Dipo- plots. Together these results may indicate that *C. baileyi* males exhibit a weak phenological shift when *Dipodomys* spp. are present, but additional data and analyses are required. The breeding onset of *C. baileyi* occurred about a month earlier than female *C.baileyi* on the same plots (Fig.1).

Models show that male *C. baileyi* on Dipo+ plots were positively associated with *C. penicillatus* biomass, and negatively associated with mean temperature (lag 0) and *C. baileyi* biomass (variation explained= 21.8%, R2= 0.17; Fig. 2). Male *C. baileyi* on Dipo- plots did not share a similar sensitivity to mean temperature but showed a similar relationship to the other variables: a positive association with *C. penicillatus* biomass and a negative association with *C. baileyi* biomass (variation explained= 44.3%; R^2^= 0.42; Fig. 2). Moreover, male *C. baileyi* on both Dipo+ and Dipo- plots exhibited similarly broad distributions in slope estimates to some environmental variables (Fig. 4), and tighter ones to some biotic factors (Appendix S1: Fig. S5). While the breadth of these distributions appears to vary little between treatments, these distributions are much broader than those exhibited by *Dipodomys* spp.

### Female desert pocket mouse (C. penicillatus)

The breeding timing of female *C. penicillatus* on Dipo+ and Dipo- plots did not significantly vary (*s*(difference=0.00, *p*= 0.73; Appendix S1: Fig. S2). However, breeding intensity was significantly higher on Dipo- than on Dipo+ plots (0.15, *p*= 0.02, deviance explained= 26.6%; Fig. 3). Thus, in the absence of *Dipodomys* spp., female *C. penicillatus* are more likely to go into reproductive mode than their counterparts on Dipo+ plots.

Female *C. penicillatus* on Dipo+ plots were positively associated with mean temperature (lag 0), warm precipitation (lag 0), cool precipitation (lag 1), and *C. penicillatus* biomass, and negatively associated with cool precipitation (lag 0) and *C. baileyi* biomass (variation explained= 55.5%; R^2^= 0.53; Fig. 2). Female *C. penicillatus* on Dipo- plots responded similarly to mean temperature (lag 0), warm precipitation (lag 0), and *C. baileyi* biomass on Dipo+ plots, but, additionally, were positively associated with NDVI (lag 1; Fig. 2). Females on both plot types exhibited relatively broad distributions in their slope estimates for environmental factors (Fig. 4). Additionally, there was only marginal overlap between *Dipodomys* and *C. penicillatus* responses to temperature and precipitation, which may suggest differences in how these environmental cues are being viewed by both species.

### Male desert pocket mouse (C. penicillatus)

Like females, the breeding timing of male *C. penicillatus* on Dipo+ and Dipo- plots did not significantly vary (*s*(difference)= 0.00, *p*= 0.44, deviance explained= 52.1%; Appendix S1: Fig. S2). Moreover, the proportion of breeding males on Dipo+ and Dipo- plots also did not significantly differ. However, despite the lack of difference in seasonal signals and breeding intensities, we noted subtle differences in the duration of breeding activity on both plots. The estimated periods of increased reproductive activity were much longer on Dipo+ than Dipo- plots (Fig. 3), which may suggest that male *C. penicillatus* prolong their breeding activities in the presence of a dominant competitor. However, without a significant treatment effect, this result should be treated cautiously.

The proportions of breeding male *C. penicillatus* on *Dipodomys* inaccessible plots were negatively associated with warm precipitation (lag 1; variation explained= 52.1%; R2= 0.48; Fig. 3) but in general males exhibited little relationship with most drivers (Dipo+ model variation explained= 63.6%; R^2^= 0.68; Fig. 2). Like male *C. baileyi*, male *C. penicillatus* on both plots also exhibited broader distributions of slope estimates to environmental variables and a tighter response to biotic factors (Fig. 4 and Appendix S1: Fig S2). Finally, male *C. penicillatus* responses to most factors did not differ from the responses of *Dipodomys* spp. (Fig. 4).

## DISCUSSION

These results show that the presence or absence of a dominant competitor can alter the reproductive phenology of small mammals by shifting the timing of the breeding season or influencing the proportion of individuals entering reproductive condition. The earlier breeding onset of *C. baileyi* females is consistent with Carter & Rudolf’s results (2022), where competitors that time their activities before or coincident with their main competitor fare better in their reproductive activities. The precise mechanism for how *Dipodomys* is influencing the timing of *C. baileyi* female phenology is unclear and we cannot distinguish between resource depletion, behavioral interference, or other *Dipodomys* spp. mediated impacts, nor can we assess whether shifts in phenology are due to individual plasticity (i.e., individuals changing the timing of their breeding activities) or selection (i.e., the presence of *Dipodomys* spp. favors individuals that would always breed early). While phenological shifts are one possible response to changes in the competitive landscape, our results demonstrate that it is not necessarily the only way species may respond. Female *C. penicillatus* exhibited no significant differences in their phenology between treatments, but a higher proportion of females entered reproductive condition in Dipo- plots. One possible explanation for *C. penicillatus*’ lower breeding intensity in Dipo+plots could be decreased body condition when *Dipodomys* spp. are present, with lower body condition potentially preventing reproduction. We do see a small, but significant (t-test p= 0.005) difference in mass between reproductive *C. penicillatus* females on Dipo+ (μ= 19.07 g) and Dipo- plots (μ= 19.75 g). Iteroparous species- i.e., with multiple opportunities to reproduce-can skip reproductive events or terminate in-process reproduction under extremely stressful environments (Stearns 1992). Thus, *C. penicillatus* females may be opting to skip a particular breeding season to wait for better conditions or be incapable of entering reproductive condition due to their own nutritional state. While Carter & Rudolf’s model (2022) does incorporate intraspecific variation, it provides limited insights for the impacts of individual-level flexibility in entering a phenological event. The results from both *C. baileyi* and *C. penicillatus* females suggest that further assessment of individual-level plasticity - in addition to intraspecific trait variation - may be useful.

Within-species phenological synchrony, which occurs when there is low variation in the timing of a biological event for a given species, may arise from consistent responses among individuals to external cues (i.e., shared responses to environmental factors at a given time scale). Carter and Rudolf (2022) posit that the benefits of low and high phenological synchrony are dependent on competitive context - specifically on the relative timing and synchrony in competing species. However, given the nature of our data, it is not immediately clear how to quantify the strength of intraspecific synchrony for our populations. If intraspecific synchrony is driven by individuals exhibiting strong and consistent responses to drivers, then tight distributions of environmental covariate estimates may indicate higher synchrony, and broader distributions may imply more coordinated responses to the environment. Consistent with the impression of higher synchrony, female *Dipodomys* spp. exhibit strong responses to environmental variables with narrower distribution curves for parameter estimates, and thus, low within-species variability in their environmental responses (Fig 4). In contrast, *Chaetodipus* spp. exhibited a broader range of responses to external stressors. This larger variation in the parameter estimates was generally consistent on both treatments and across sexes, suggesting that high intraspecific variability in these species may not be driven solely by the direct impacts of competition with *Dipodomys* spp. This may mean that these *Chaetodipus* spp. exploit environmental conditions that are underutilized by *Dipodomys* spp. as an adaptive strategy.

While shifts in biotic context appear to drive changes in the breeding patterns of the females for both *Chaetodipus* spp., it did not appear to be the case for males. The sex-related differences in response to environment and competitive landscape are not necessarily surprising since, in mammals, female reproduction is often limited by resource availability for gestation and lactation and male reproduction by access to mates (Mysterud et al 2008). Because female mammals are energetically and nutritionally providing for their own increased resource needs and those of their offspring, females may be particularly sensitive to the effects of resource depletion caused by interspecific competition. This may explain the observed phenological shifts of female *C. baileyi* and increased breeding intensity of *C. penicillatus* in Dipo- plots. In contrast, males for many mammal species - including those at our field site - are frequently uninvolved in offspring rearing. This may explain the weak and lack of seasonal signal for breeding detected in male *C. baileyi* and *C. penicillatus*, respectively. If male energetic costs are primarily due to acquiring and defending females from other conspecifics (Ostfeld 1985), then their phenology may be less sensitive to interspecific competition and more driven by female reproductive changes and intraspecific competition for mates.

Our results also show that the effects of competition on phenology may be species specific. While both *Chaetodipus* spp. have repeatedly demonstrated that *Dipodomys* have strong negative impacts on their population sizes (Brown and Munger 1984, Ernest and Brown 2001, Bledsoe and Ernest 2019), they showed different phenological responses to the removal of *Dipodomys*. Clearly, more empirical and theoretical research is required to better understand how and when competition will cause shifts in timing as opposed to prevalence. The differences in model fits between treatments also suggest that continual reassessment of environmental drivers of phenology may be warranted as biodiversity change alters the competitive environments many species are experiencing. Future studies that can tease apart the direct and indirect impacts of competition may be useful in developing a robust understanding of the effects of competition in species’ phenology. By recognizing that competitive landscapes can influence the responses of breeding individuals, we gain a more nuanced understanding of one of the key biological phenomena that can alter ecosystem dynamics.

## ACKNOWLEDGMENTS

We thank the numerous volunteers, staff, and research assistants, who collected the Portal Project data over the years. This work was supported by research grants awarded to SKME by the US National Science Foundation (NSF DEB-1929730) and USDA National Institute of Food and Agriculture, Hatch Project FLA-WEC-005983.

