## Appendix for "Shifts in competitive structures can drive variation in species’ phenology"

**Journal:** Ecology

**Manuscript type:** Article

**Open Research Statement:** Data (10.5281/zenodo.7301687) and code (10.5281/zenodo.8125113) used to conduct data analyses are archived in Zenodo.

**Key words**: breeding phenology, competition, generalized additive model (GAM), phenological synchrony, population dynamics, species interactions

### APPENDIX S1

#### Methods

*Breeding phenology of C. penicillatus under different C. baileyi regimes*

To account for the known changes in the *C. baileyi* population over time, which may have influenced *C. penicillatus* population dynamics, we also explored the breeding phenology of *C. penicillatus* and their drivers during three different periods: 1977-1994 (pre-*C. baileyi* establishment), 1995-2010 (*C. baileyi* establishment), 2011-2014 (post-*C. baileyi* establishment). The time span of each period was based on the results discussed by Bledsoe and Ernest (2019).

*Associations between factors and rodent reproduction under different biotic conditions*

To test the relative importance of different abiotic and biotic factors in driving reproductive phenology, we built models of increasing complexity. The first model included only abiotic factors (NDVI, temperature, precipitation). The second model included abiotic factors and a proxy for intraspecific competition (population-specific biomass). Finally, the third model included abiotic factors and proxies for intra- and interspecific competition (biomass of competitors). We assessed model performance based on $R^2$ values and percentage of deviance explained.

#### Results

*Breeding phenology of C. penicillatus under different C. baileyi regimes*

There was no detectable trend in the breeding proportions of male and female *C. penicillatus* in both control and exclosure plots from 1977 to 1994, a period when *C. baileyi* were also uncommon (Appendix Fig. 1a and 1d). After five years, in 1995 to 2010, when *C. baileyi* got established in the site, the breeding timing of females in both *Dipodomys* accessible (Dipo+) and inaccessible (Dipo-) plots occurred around the same time. The peak period of reproduction for females in both plots occurred around May to June (Appendix Fig. 3b). During the same period, males in Dipo- plots exhibited a relatively earlier peak reproductive period, around March, than those in Dipo+ plots. The breeding onsets of males in both plots occurred before females. Finally, in 2011 to 2014, a period when *C. baileyi* were again less common, the breeding proportions of both males and females were lower compared to the two other regimes (Appendix Fig. 3c and 3f). Females in Dipo- plots exhibited a more pronounced peak reproductive period, in May, compared to those in Dipo+ plots (Appendix Fig 3c). On the other hand, males in control plots exhibited a more pronounced peak reproductive period (Appendix Fig.3f). The disappearance of timing differences for *C. penicillatus* on Dipo+ and Dipo- plots when *C. baileyi* is also prevalent could indicate that the lack of phenology change reported in the main text is because *C. penicillatus* did not perceive a difference in the competitive environment on Dipo+ and Dipo- plots. Strong competition between *C. penicillatus* and *C. baileyi* has been demonstrated previously (Bledsoe and Ernest 2019) and more investigation of this dynamic may prove informative.

*Associations between factors and rodent reproduction under different biotic conditions*

Across all species, sexes, and treatments, models that included both abiotic and biotic factors performed relatively better than simpler models (Appendix Table 1). Additionally, the breeding proportions of male and female *C. penicillatus* during different *C. baileyi* regimes were also best fit using complex models that included all abiotic and biotic factors (Appendix Table 2). However, there was no predictor that was consistently important across all models

(Appendix Fig 4). In most cases, influential abiotic predictors were those that were not lagged values.

Responses of females of both *Chaetodipus* spp. to various abiotic and biotic factors were relatively more disparate from *Dipodomys* spp. than males in Dipo+ plots (i.e., less overlap in coefficient estimates for different covariates; Appendix Fig. 5, 6) based on the overlap in the distribution of coefficient estimates. Additionally, the range of coefficient estimates were much broader for females. This may suggest a more diverse relationship between proportion of breeding females in control plots and covariates compared to males. Similarly, females of both *Chaetodipus* spp. in Dipo- plots exhibited more disparate responses to covariates with each other compared to males (Appendix Fig. 5, 6).

**Appendix Tables**

**Appendix S1: Table S1.** Performance of models (% deviance explained) with increasing levels of complexity built to explain the variation in the proportion of breeding male and female *Chaetodipus baileyi* and *C. penicillatus* in *Dipodomys* accessible (Dipo+) and inaccessible (Dipo-) plots in a long-term monitoring site in Portal, Arizona from 1977 to 2014. Values enclosed in parentheses are $R^2$.

| *C. baileyi* | | | | |
|---|---|---|---|---|
| Model: | male | | female | |
| | Dipo+ | Dipo- | Dipo+ | Dipo- |
| Abiotic factors only | 18.2 (0.11) | 38.1 (0.37) | 66.9 (0.56) | 66.7 (0.65) |
| Abiotic + intraspecific competition | 18.5 (0.10) | 40.8 (0.39) | 66.2 (0.57) | 66.7 (0.65) |
| Abiotic + intra- + interspecific competition | 21.8 (0.17) | 44.3 (0.42) | 66.4 (0.57) | 69.3 (0.66) |
| *C. penicillatus* | | | | |
| Model: | male | | female | |
| | Dipo+ | Dipo- | Dipo+ | Dipo- |
| Abiotic factors only | 62.2 (0.64) | 51.9 (0.49) | 52. 5 (0.50) | 46.6 (0.40) |
| Abiotic + intraspecific competition | 62.4 (0.65) | 52.0 (0.49) | 50.6 (0.49) | 47.6 (0.41) |
| Abiotic + intra- + interspecific competition | 63.6 (0.68) | 52.1 (0.48) | 55.5 (0.53) | 50.4 (0.44) |

**Appendix S1: Table S2**. Performance of models used to fit data on breeding proportions of female and male *C. penicillatus* in *Dipodomys* accessible (Dipo+) and inaccessible (Dipo-) plots during different *C. baileyi* regimes based on percent (%) deviance explained. In all models, year was assumed to have a linear effect and not added as a smooth term to avoid overfitting.

| Model | **1977-1994** | | **1995-2010** | | **2011-2014** | |
|---|---|---|---|---|---|---|
| | **Dipo+** | **Dipo-** | **Dipo+** | **Dipo-** | **Dipo+** | **Dipo-** |
| **Female** | | | | | | |
| Abiotic only | NA | NA | 61.8 | 45.7 | 78.6 | 63.6 |
| Abiotic + intraspecific competition | NA | NA | 62.4 | 45.7 | 82.2 | 67.0 |
| Abiotic + intra- + interspecific competition | NA | NA | 64.9 | 47.6 | 92.8 | 79.0 |
| **Male** | | | | | | |
| Abiotic only | NA | NA | 61.7 | 62.1 | 80.2 | 65.3 |
| Abiotic + intraspecific competition | NA | NA | 63.4 | 62.1 | 80.2 | 75.4 |
| Abiotic + intra- + interspecific competition | NA | NA | 65.7 | 62.1 | 82.4 | 82.0 |

**Appendix Figures**

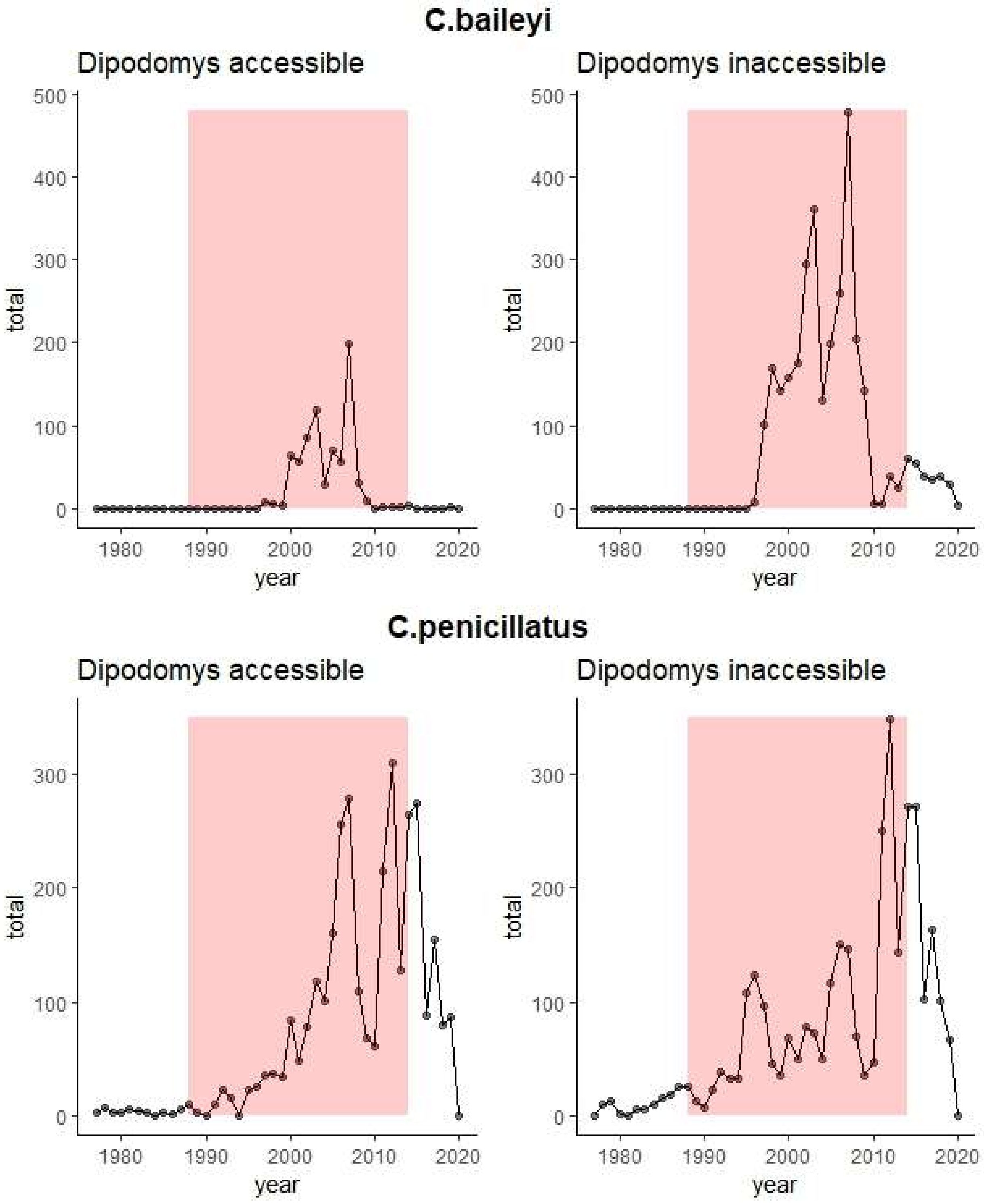

**Appendix Fig. S1.** Abundances of *C. baileyi* (top panel) and *C. penicillatus* (bottom panel) at an experimental field site near Portal, AZ from 1977-2020. Period highlighted in red (1988-2014) was the period used in the analyses of this study.

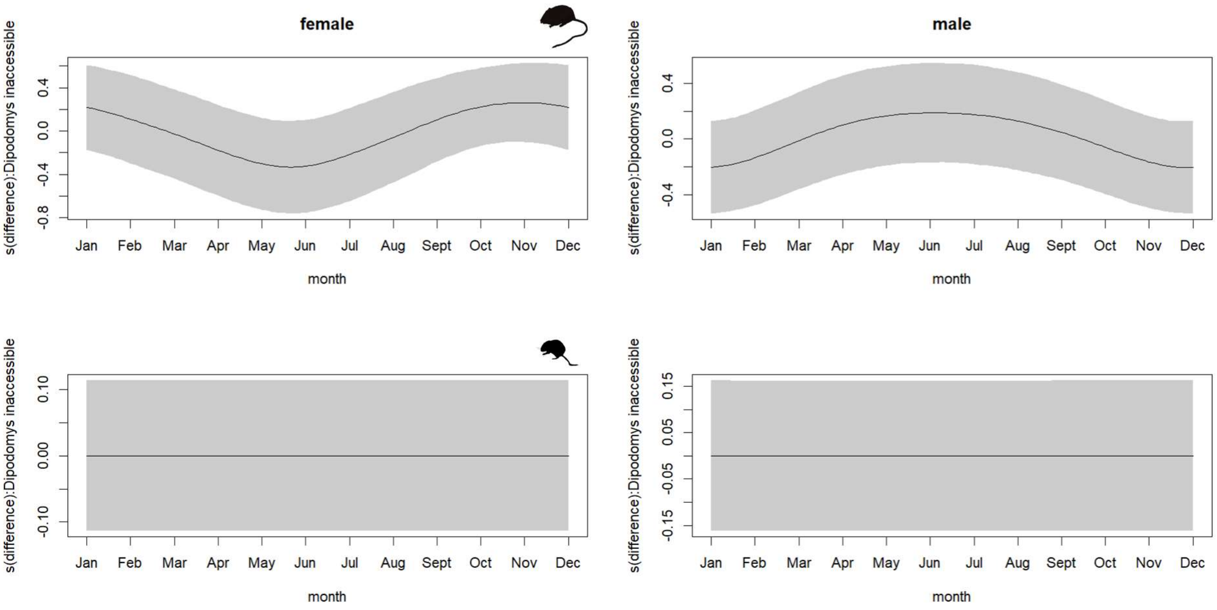


**Appendix Fig. S2.** Estimated differences in the seasonal trends of female (left panel) and male (right panel) *C. baileyi* (top panel) and *C. penicillatus* (bottom panel) on *Dipodomys* accessible and *Dipodomys* inaccessible plots.

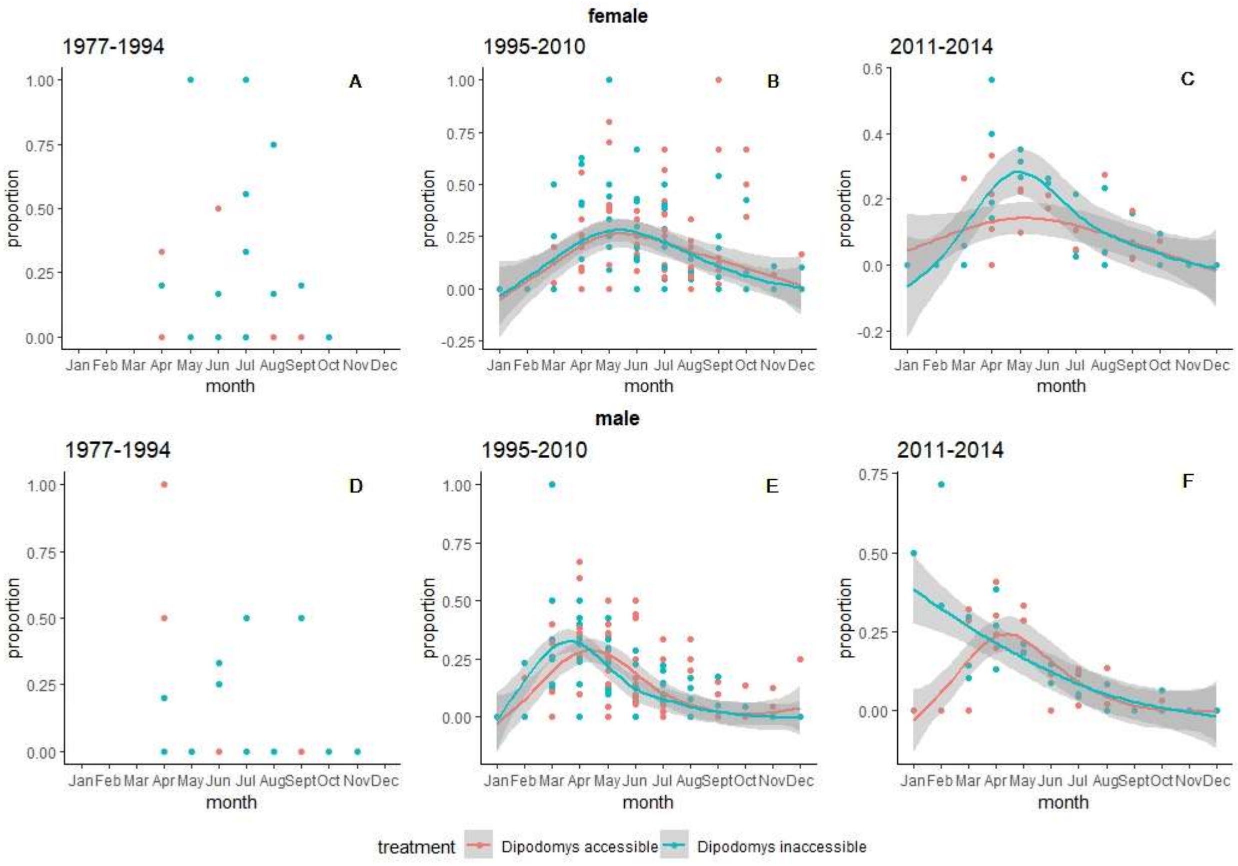


**Appendix Fig. S3.** Breeding phenology of female (top panel) and male (bottom panel) desert *C. penicillatus* in *Dipodomys* accessible and inaccessible plots during three different periods of *C. baileyi* establishment in a long-term experimental site near Portal, Arizona. Each filled circle represents raw data. Trend lines were estimated using generalized additive models.

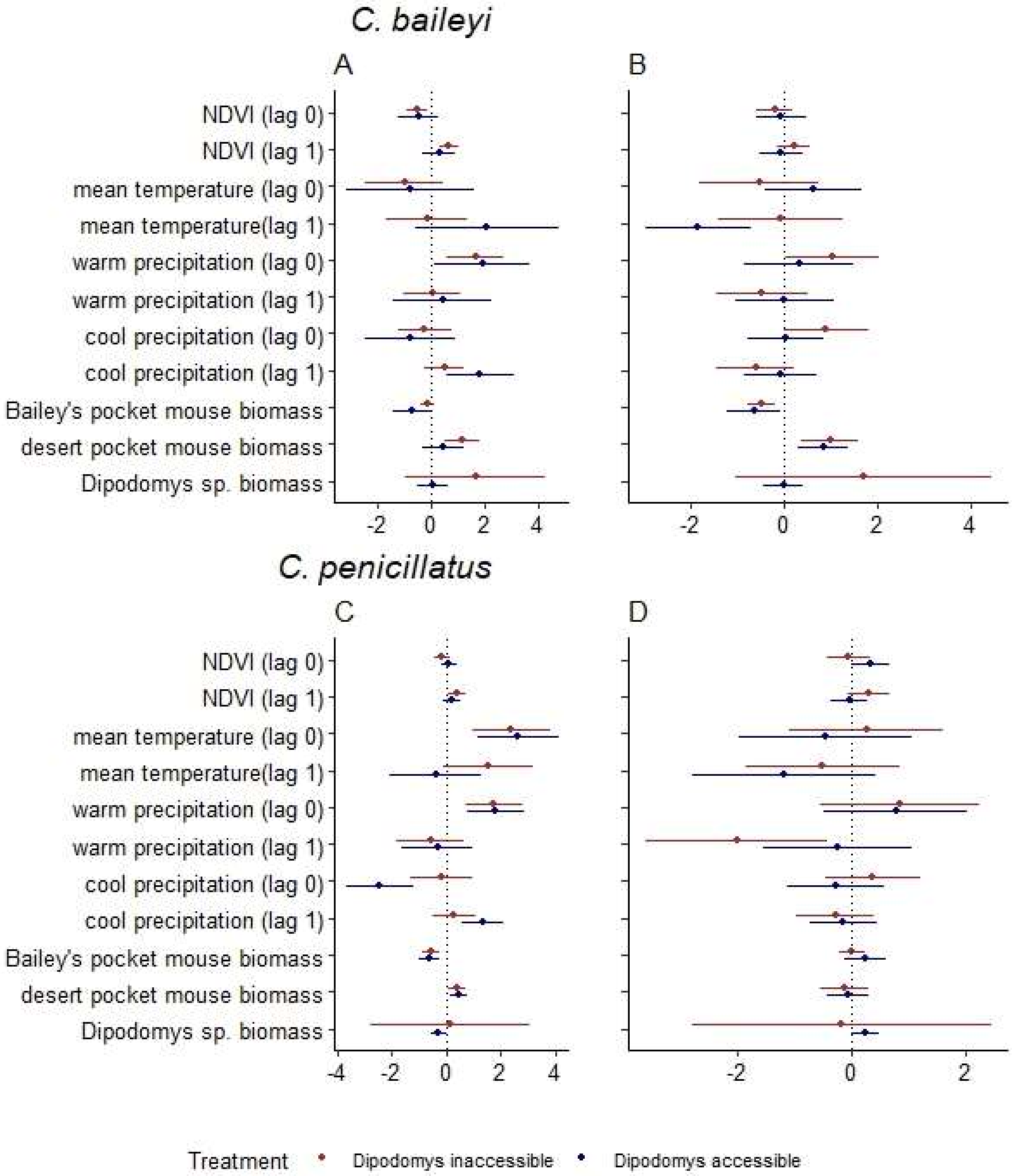


**Appendix Fig. S4**. Association between abiotic and biotic factors with the proportion of reproductive female (A and C) and male (B and D) *C. baileyi* (top panel) and *C. penicillatus* (bottom panel). Filled circles indicate the mean, and lines indicate the 95% confidence interval of the coefficient estimates. Estimates were derived from the generalized additive models (GAMs) fit for each sex- and treatment-specific dataset.

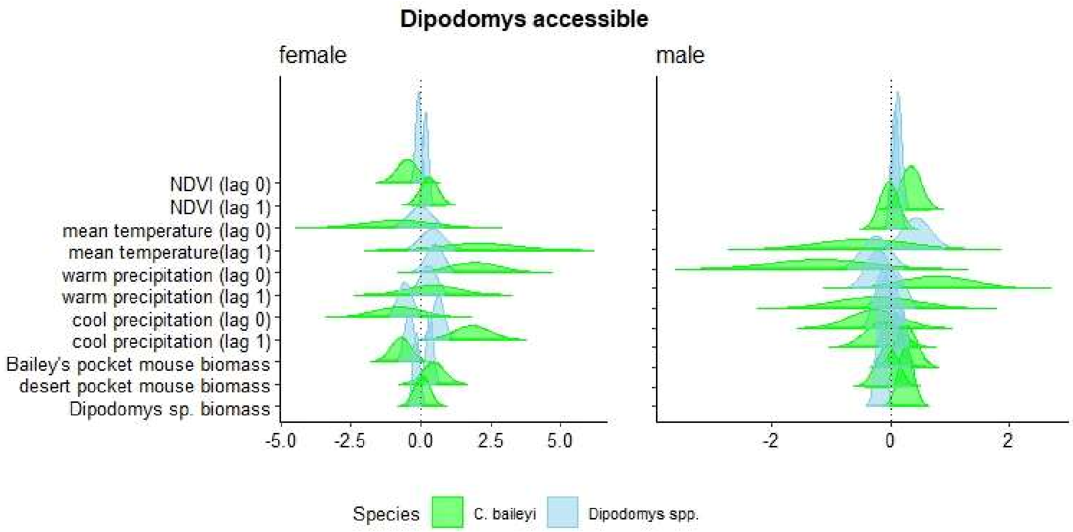
Dipodomys accessible
female
male
NDVI (lag 0)
NDVI (lag 1)
mean temperature (lag 0)
mean temperature(lag 1)
warm precipitation (lag 0)
warm precipitation (lag 1)
cool precipitation (lag 0)
cool precipitation (lag 1)
Bailey's pocket mouse biomass
desert pocket mouse biomass
Dipodomys sp. biomass
-5.0
-2.5
0.0
2.5
5.0
-2
0
2
Species
C. baileyi
Dipodomys spp.

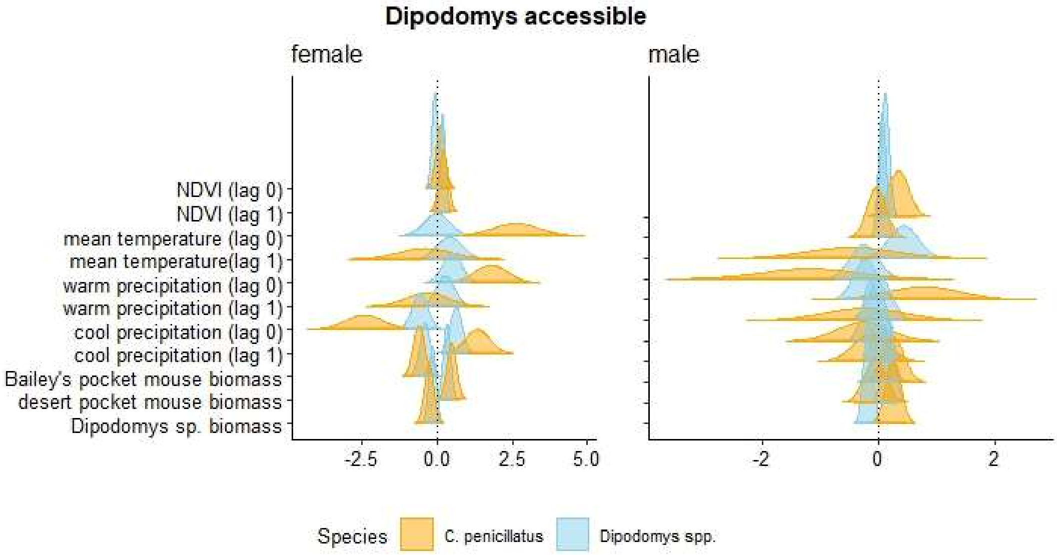
Dipodomys accessible
female
male
NDVI (lag 0)
NDVI (lag 1)
mean temperature (lag 0)
mean temperature(lag 1)
warm precipitation (lag 0)
warm precipitation (lag 1)
cool precipitation (lag 0)
cool precipitation (lag 1)
Bailey's pocket mouse biomass
desert pocket mouse biomass
Dipodomys sp. biomass
-2.5
0.0
2.5
5.0
-2
0
2
Species
C. penicillatus
Dipodomys spp.

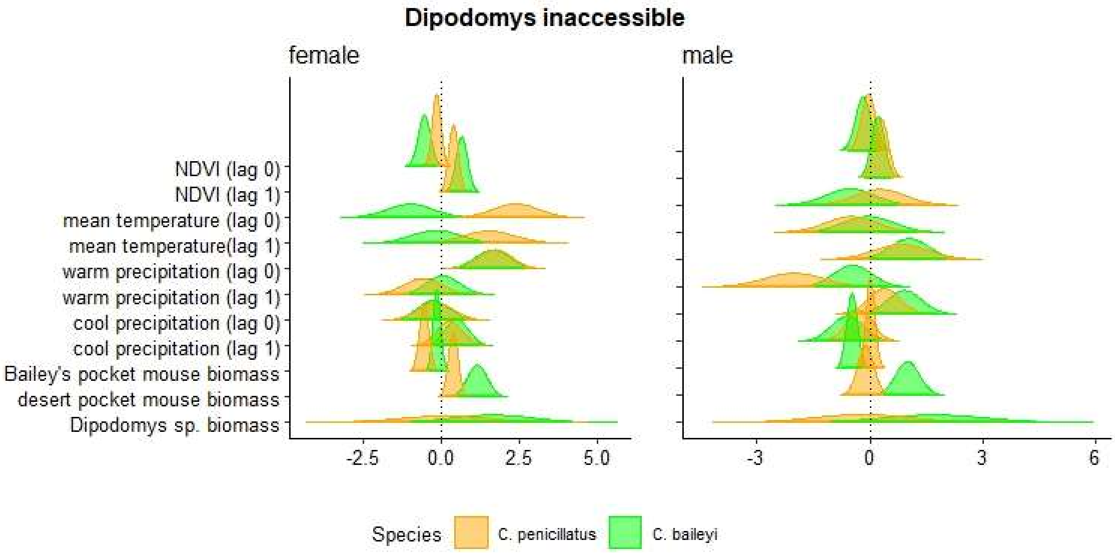
Dipodomys inaccessible
female
male
NDVI (lag 0)
NDVI (lag 1)
mean temperature (lag 0)
mean temperature(lag 1)
warm precipitation (lag 0)
warm precipitation (lag 1)
cool precipitation (lag 0)
cool precipitation (lag 1)
Bailey's pocket mouse biomass
desert pocket mouse biomass
Dipodomys sp. biomass
-2.5
0.0
2.5
5.0
-3
0
3
6
Species
C. penicillatus
C. baileyi

**Appendix Fig. S5.** Association between all abiotic and biotic factors with the proportion of reproductive female (left panel) and male (right panel) *C. baileyi* (top panel) and *C. penicillatus* (middle panel) relative to *Dipodomys* spp.in *Dipodomys* accessible plots and relative to each other in *Dipodomys* inaccessible plots (bottom panel). Distribution of coefficient estimates were derived from the generalized additive models (GAMs) fit for each sex and treatment.

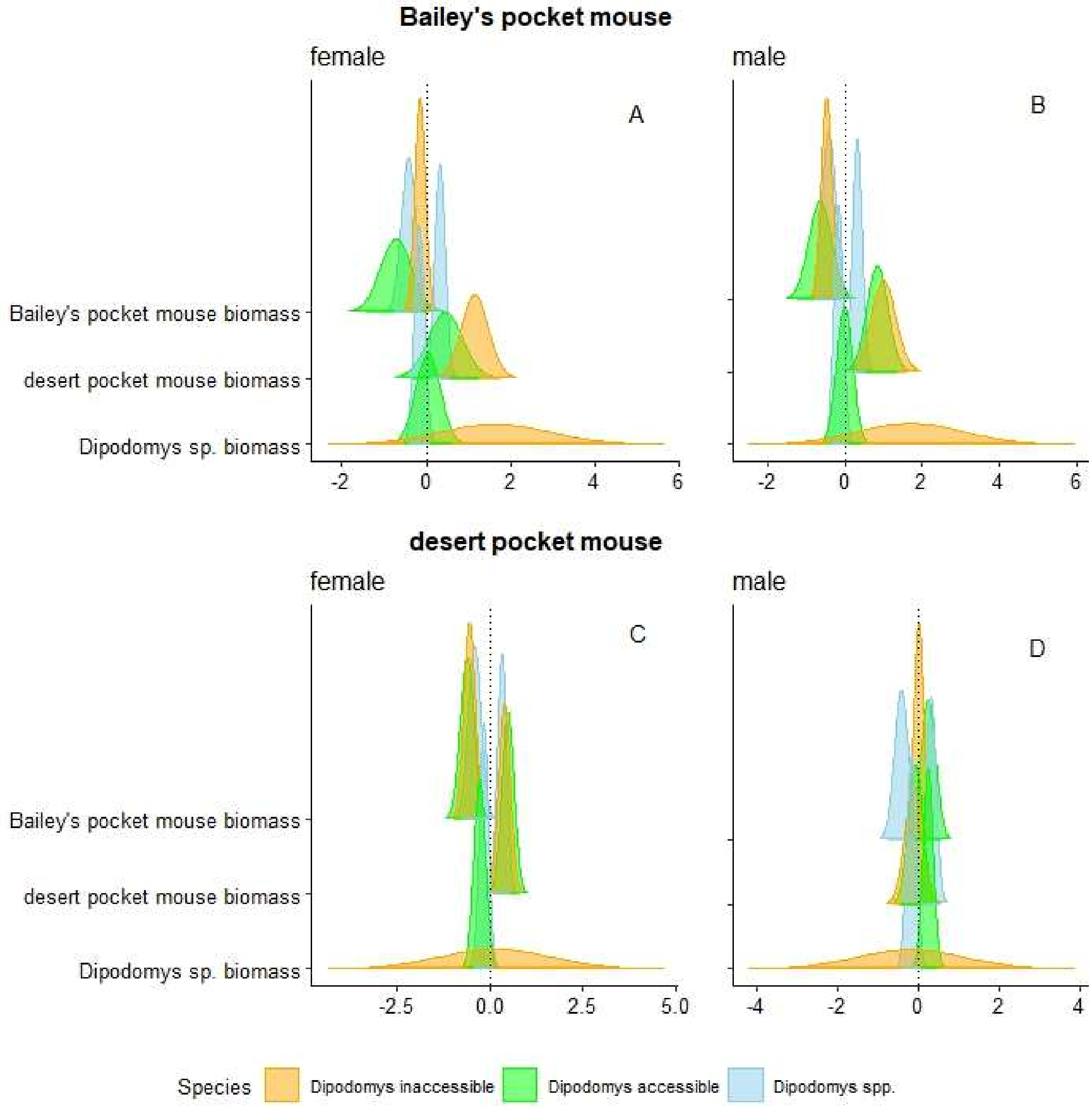


**Appendix Fig. S6.** Association between select biotic factors with the proportion of reproductive female (left panel) and male (right panel) *C.* baileyi (top panel) and *C. penicillatus* (bottom panel) in *Dipodomys* accessible and inaccessible plots relative to *Dipodomys* spp.. Distribution curves indicate the normal distribution with mean equal to the point estimate, and standard deviation equal to the standard error of the coefficients. Estimates were derived from the generalized additive models (GAMs) fit for each sex- and treatment-specific dataset.
